# The potential role of collagens in congenital Zika syndrome: A systems biology approach

**DOI:** 10.1101/541268

**Authors:** Renato S Aguiar, Fabio Pohl, Guilherme L Morais, Fabio CS Nogueira, Joseane B Carvalho, Letícia Guida, Luis WP Arge, Adriana Melo, Maria EL Moreira, Daniela P Cunha, Leonardo Gomes, Elyzabeth A Portari, Erika Velasquez, Rafael D Melani, Paula Pezzuto, Fernanda L de Castro, Victor EV Geddes, Alexandra L Gerber, Girlene S Azevedo, Bruno L Schamber-Reis, Alessandro L Gonçalves, Inácio Junqueira-de-Azevedo, Milton Y Nishiyama, Paulo L Ho, Alessandra S Schanoski, Viviane Schuch, Amilcar Tanuri, Leila Chimelli, Zilton FM Vasconcelos, Gilberto B Domont, Ana TR Vasconcelos, Helder I Nakaya

## Abstract

Zika virus (ZIKV) infection during pregnancy could cause a set of severe abnormalities in the fetus known as congenital Zika syndrome (CZS). Experiments using animal models and *in vitro* systems significantly contributed to our understanding of the physiopathology of ZIKV infection. However, the molecular basis of CZS is not yet studied in humans. Here, we used a systems biology approach to integrate transcriptomic, proteomic and genomic data from post-mortem brains of neonates with CZS. We observed that collagen genes were greatly reduced in CZS brains at both the RNA and protein levels and that neonates with CZS have several polymorphisms in collagen genes associated with osteogenesis imperfect and arthrogryposis. These findings were validated using immunohistochemistry and collagen staining of ZIKV infected and non-infected samples. Additionally, it was found that cell adhesion genes that are essential for neurite outgrowth and axon guidance were up-regulated and thereby confirmed the neuronal migration defects observed. This work provided new insights into the underlying mechanisms of CZS and revealed host genes associated with CZS susceptibility.

## Introduction

Zika virus (ZIKV) infection during pregnancy is associated with several neurological problems in the fetus^1,2^. Collectively, the set of abnormalities is known as congenital Zika syndrome (CZS) and could involve microcephaly, brain calcifications, ventriculomegaly, cortical malformations due to migration disorders including agyria/lissencephaly, congenital contractures, and ocular abnormalities^1,3,4^. In adults, the most common symptoms of ZIKV infection are fever, rash, arthralgia, conjunctivitis, and headache^5^. Although most pregnant women exposed to ZIKV give birth to healthy babies, 0.3–15% of cases develop CZS^6^. The frequency of infant deaths (miscarriages and perinatal deaths) is low (∼1% of CZS), and most of them present intrauterine akinesia syndrome (arthrogryposis)^7^.

Several studies *in vitro* and with brain organoids and neurospheres demonstrated that ZIKV could directly infect human neural progenitor cells^8^, impairs the cortical development^9^, affects neuron migration impacting brain size^10^, and promotes brain malformation^11^. Nevertheless, the molecular basis of CZS and susceptibility genes associated with the most severe cases in human newborns remains unknown.

Systems biology approaches were successfully applied to reveal the molecular mechanisms associated with viral infection and vaccination^12,13^. By integrating different types of omics data, systems biology provides a global overview of the network of genes, transcripts, proteins, and metabolites involved with a biological condition or perturbation^14^. When applied to human infectious diseases, it could provide critical insights into the complex interplay between pathogen and host, thereby leading to potential novel intervention strategies.

In this study, we have generated genomic, transcriptomic and proteomic data from the blood and post-mortem brain samples of eight neonates with confirmed ZIKV infection during pregnancy and with no congenital genetic diseases nor another STORCH group vertical transmission. After three-layer omics data integration, we highlighted the molecular pathways underlying neurological damage. Systems biology combined with histopathological analysis revealed that genes associated with matrix organization were dramatically down-regulated in the brain of neonates with CZS, which could explain the neuronal migration disorders and microcephaly attributed to ZIKV infection.

## Results

### Neonates with severe CZS

From October 2015 to July 2016 we followed a group of pregnant women with symptoms of ZIKV exposition at distinct weeks of gestation and from two endemic areas in Brazil—northeast (Campina Grande, Paraiba state) and southeast (Rio de Janeiro, Rio de Janeiro state) regions. During this period, we enrolled pregnant women who were referred to public healthcare with a history of rash or fetus with central nervous system (CNS) abnormality confirmed by ultrasonography or magnetic resonance imaging, as well as postnatal physical examination suggestive of microcephaly.

We focused on eight neonates that had died in the first 48 hours postpartum with severe arthrogryposis (Figure 1a). ZIKV genome was detected in all cases during pregnancy by RT-PCR in clinical samples from mothers and the neonates such as urine, plasma, amniotic fluid, placenta, and umbilical cord. We also detected the virus genome through RT-PCR and *in situ* hybridization in fetal post-mortem tissues (Figure 1b). Other microcephaly causes including congenital genetic diseases, infection with arboviruses that circulate in the same area (Dengue and Chikungunya) and teratogenic pathogens (STORCH) were all excluded (Table S1).

**Figure 1.**
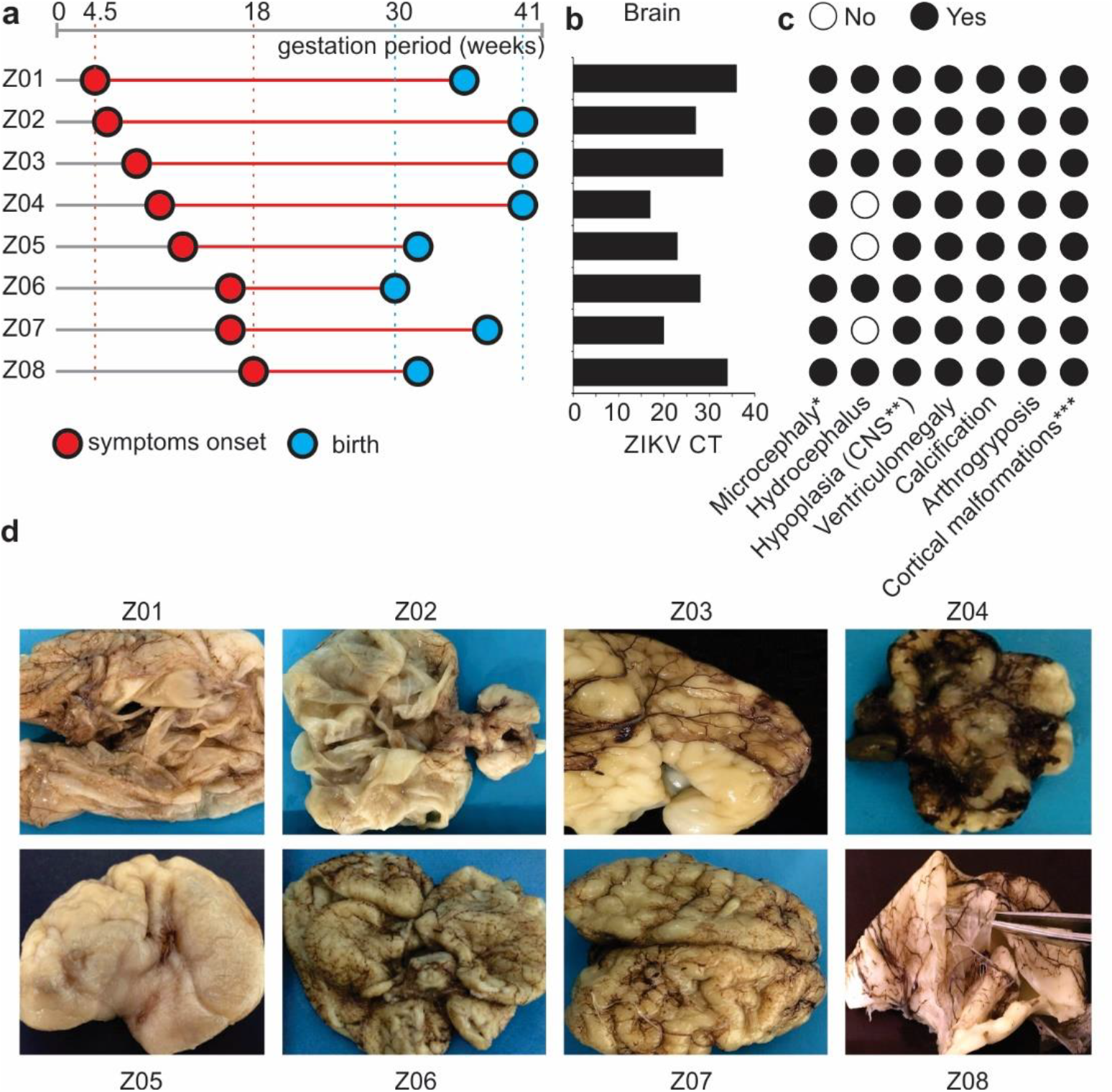
Clinical diagnoses and brain damage of deceased neonates with CZS. (a) Gestation timeline for the eight neonates with CZS. The symptoms include fever, exanthema, arthralgia, conjunctivitis, and headache in pregnant women during gestation. (b) Zika genome detection by RT-PCR from post-mortem brain samples expressed in CT values. (c) Lesions in the central nervous system (CNS) of neonates with CZS investigated by prenatal ultrasound and MRI examinations; *At birth only 3 cases (Z04,Z05, Z07) had microcephaly. The others had normal or enlarged cephalic perimeter due to obstructive hydrocephalus; **Cerebellum and Brainstem hypoplasia; ***cortical malformations due to neuronal migration disturbance (agyria, polymicrogyria or lissencephaly) (d) Brains from autopsies showing various degrees of lesions, including collapse due to hydrocephalus and small brains with few gyri or agyria and consistently with severe loss of CNS structures and congested leptomeninges. The numbers of Zika cases are depicted as presented in Table S1.

Five out of eight cases of CSZ showed ZIKV exposition symptoms in the first trimester of pregnancy corroborating with other reports^15,16^ that describe increasing risk of microcephaly at the beginning of gestation (Figure 1a). Microcephaly was observed in the early gestation weeks through ultrasonography in all cases. However, the cephalic perimeter at birth was considered normal (higher than 32 cm) in most of the neonates due to severe ventriculomegaly/obstructive hydrocephalus. The brain usually collapsed after removal of the skull during autopsy showing tiny brains in all cases (on average 66 grams; ranging from 7 to 180 grams). A detailed neuropathological description of all cases has been previously reported^10^.

The brains with higher viral load (lower cycle threshold or CT values) exhibited the most destructive patterns of CNS structures (Figure 1b and Table S1). Macroscopic observations showed thickened and congested leptomeninges, very thin parenchyma and corpus callosum, and asymmetric ventriculomegaly (Figure 1d). Shallow sulci or agyria was prevalent in all cases (Figure 1d). The hippocampus, basal ganglia, and thalami were usually not well identified and malformed. Cerebellar hypoplasia was observed in all cases, with an irregular cortical surface and calcification foci were detected macroscopically (Figure 1d). The brainstem was deformed and hypoplastic in most of the cases.

The histopathological analysis confirmed the migration disturbances represented by abnormal immature cell clusters along the white matter and over pia mater (Table S2). An intense immune response to cell injury was observed in all cases as demonstrated by the gliosis and inflammatory infiltrate (T-lymphocytes and histiocytes) in the meninges, cerebral hemispheres, and spinal cord (Tables S1 and S2). Reduction of the descending motor fibers was also observed. The histopathological analysis also displayed a loss of motor nerve cells in the spinal cord and atrophy of the skeletal muscle. These could explain the intrauterine akinesia and consequent arthrogryposis observed in all cases (Tables S1 and S2).

### Transcriptome and Proteome analyses of CZS Brains

We utilized high-throughput sequencing and mass spectrometry technologies to assess the changes in the transcriptome and proteome of CZS brains (Z03, Z05, and Z08 in Figure 1) compared to the control brain (Edwards’syndrome). Differential expression analysis revealed 509 genes associated with CZS, of which 228 were up-regulated and 281 were down-regulated in ZIKV-infected neonates (Figure S1a and Table S3). Among the pathways enriched with up-regulated genes, we found the “Unblocking of NMDA Receptor, Glutamate Binding, and Activation” and “Glutamate Neurotransmitter Release Cycle” (Figure S1a and Table S3). These findings support our previous *in vitro* work showing that the blockage of the NMDA receptor prevents the neuronal death induced by ZIKV infection^17^. Among the pathways enriched with down-regulated genes, we found collagen formation, glucose metabolism, signaling by TGF-beta receptor complex, Class I MHC mediated antigen processing and presentation, and amyloid fiber formation (Figure S1b). These down-regulated genes indicate that ZIKV infection could affect immune-response pathways, cellular metabolism and the very formation of connective tissue in the brain.

Cell adhesion genes—essential for neuronal migration and recruiting of immune cells including NCAM receptors—are up-regulated in the CZS brains, which corroborates the migration disturbance and inflammatory infiltration events (CD8+ T-lymphocytes and CD68+ histiocytes) observed in the histopathological analysis (Table S3). Collagen genes (*COL1A1, COL1A2, COL3A1, COL5A1, COL5A2, COL6A3, COL12A1*, and *COL14A1*) essential for the development of the brain and the blood-brain barrier^18^ are down-regulated in the CZS brains.

We subsequently investigated the protein levels in CZS brains compared to the ZIKV negative control. The proteomic analysis identified 252 and 110 proteins up- and down-regulated in CZS brain respectively (Figure 2a and Table S4). Furthermore, a set of proteins were exclusively detected either in brains with CZS (714 proteins) or in the ZIKV negative control brain (79 proteins) (Figure 2a and Table S4). Similar to the transcriptomic analysis, up-regulated proteins were enriched for “glucose metabolism” and “L1CAM interactions,” whereas down-regulated proteins were enriched for “extracellular matrix organization” and “collagen formation” (Figure 2b and Table S4). Among the proteins down-regulated in all three neonates with CZS compared to control, COL1A1, COL1A2, PPIB, SERPINH1, and OGN were found. While PPIB is instrumental in collagen trimerization, SERPINH1 is critical to collagen biosynthesis^19^. Additionally, the functions of OGN in the extracellular matrix are related to collagen fibrillogenesis, cell proliferation, and development, as well as osteoblast differentiation and bone development^20^.

**Figure 2.**
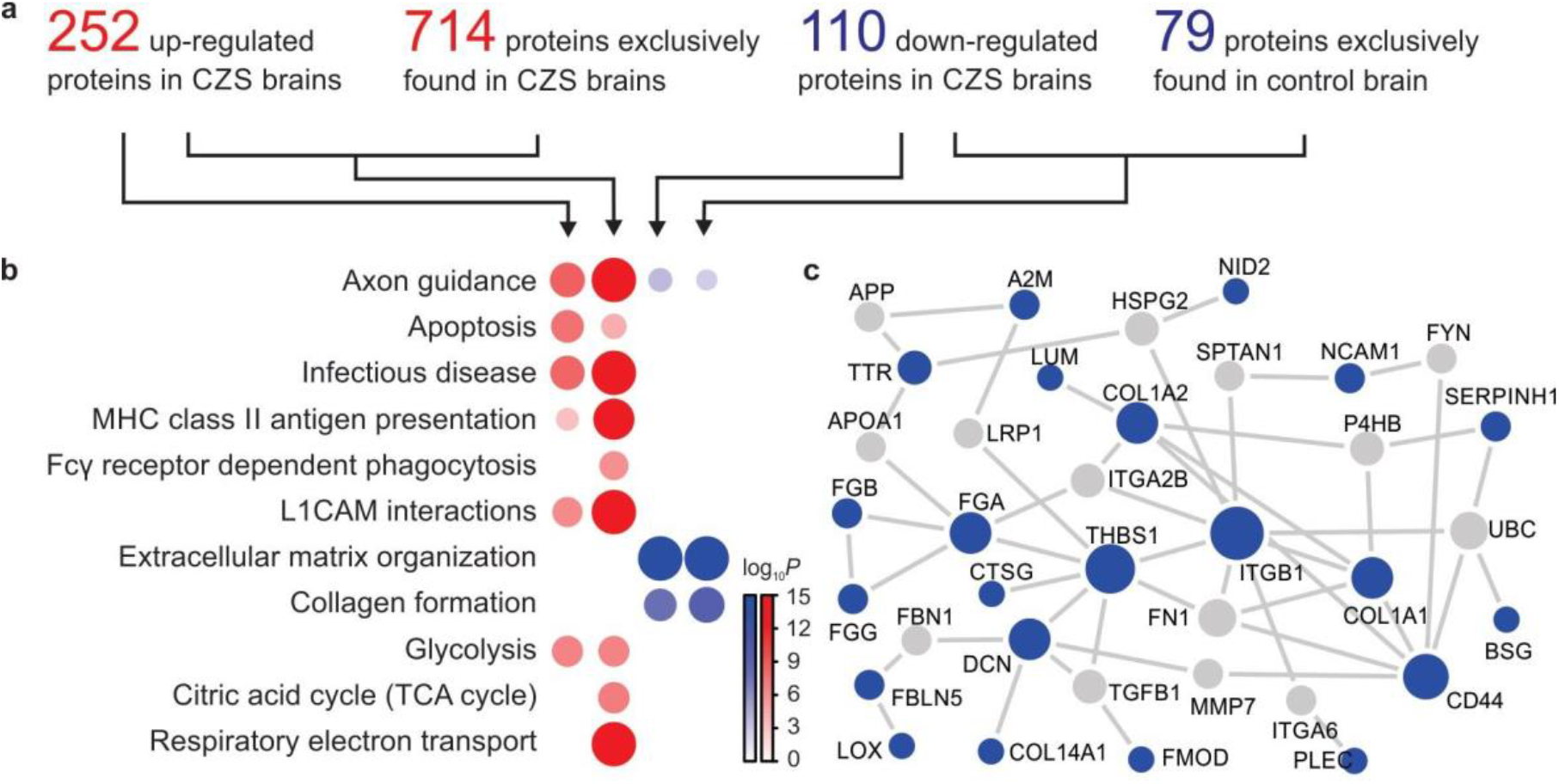
Proteins related to CZS and microcephaly. (a) The number of differentially expressed proteins up- (red) and down-regulated (blue). Proteins with adjusted p-value < 0.05 were considered differentially expressed. Proteins expressed only in one condition were considered exclusively expressed proteins. (b) Enrichment of functional pathways for proteins found in CZS brains. Red represents the up-regulated proteins, and blue represents the down-regulated proteins. Adjusted p-value (-log10) of Over Representation Analysis is indicated by color intensity and circle size. (c) Protein-protein interactions for down-regulated proteins. Blue nodes indicate the down-regulated proteins observed and grey nodes indicate the additional proteins. The circle size represents the node degree.

Protein-protein interaction data obtained from the STRING database was used to assess the interacting proteins related to the brain’s connective tissue (Figure 2c). The interactome showed the same pattern of the down-regulation of essential proteins hubs involved in collagen formation (COL1A1 and COL1A2) and adhesive glycoprotein that mediates cell-to-cell and cell-to-matrix interactions (ITGA2B, NCAM, FNB, IGB1, and THBS1). LOX is down-regulated in both transcriptome and proteome analysis and plays a key role in cross-linking fibers of collagen and elastin. The down-regulation of collagen pathways in the brain endothelia could partially explain the vascular problems and ischemia events observed in CZS neonates. Once again, proteomics analysis validates transcriptomics as well as the macroscopic and microscopic images showing the modulation of proteins involved in brain architecture matrix and neuronal migration disorders in ZIKV affected neonates. The interactome showed the modulation of fibrinogen components (FGA, FGB, and FGG), which are components of blood clots and are formed following vascular injury. These findings relate to the intense blood congested leptomeninges found in the CSZ brains (Figure 2c and 1d).

We subsequently cross-referenced the lists of genes and proteins that were differentially expressed in CZS compared to the negative control and found, respectively, 12 and 23 up- and down-regulated shared genes and proteins (Figure 3a). The functions of several of these genes could provide insights into ZIKV neuropathogenesis. For instance, *NCAM1* is essential for neurite outgrowth, *COL1A1* and *COL1A2* genes encode the alpha 1 and 2 chains of collagen type I, and *PRDX2* regulates the antiviral activity of T cells (Figure 3a). For *TTR* and *AGT* genes, however, the levels of the RNA were higher in CZS than in control (up-regulated at RNA level) whereas the protein levels were lower in CZS (down-regulated at protein level). Similarly, eight genes were up-regulated in proteomics but down-regulated in transcriptomics. These inverted patterns between RNA and protein levels could partially be due to post-transcriptional regulation mechanisms that include miRNAs. Thus, we checked whether genes that were up-regulated in transcriptomics but not up-regulated in proteomics dataset were known miRNA targets. Our *in silico* approach predicted that eight miRNAs were induced upon infection and possibly involved regulation of genes related to CZS (Figure 3b). Among them, mir-17-5p was already shown to be induced by flavivirus infections, including ZIKV infection in astrocytes^21^.

**Figure 3.**
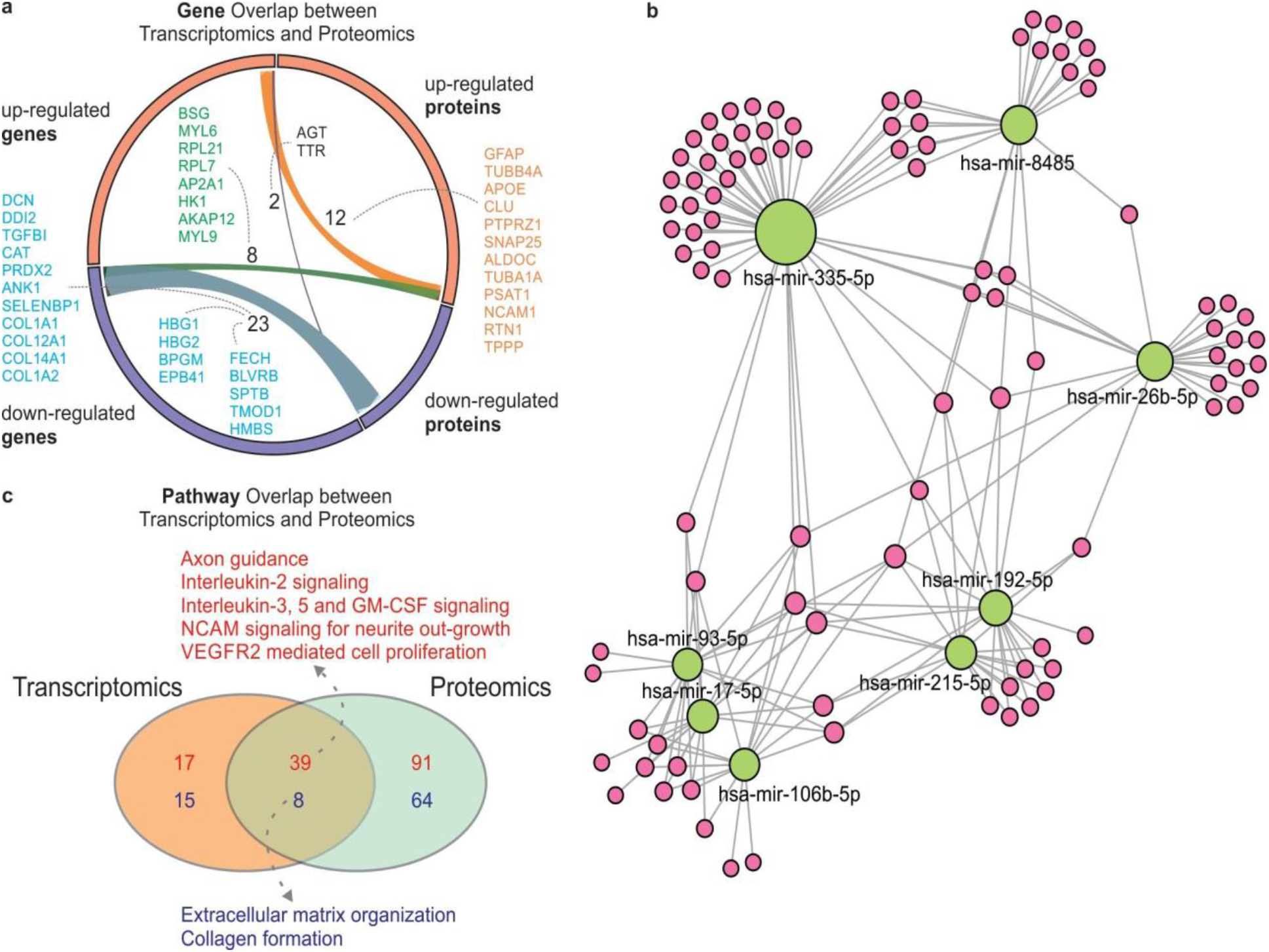
Transcriptomics and proteomics interplay in CZS. (a) The overlap between differentially expressed genes (DEGs) and differentially expressed proteins (DEPs) in CZS. The fraction of up- and down-regulated genes/proteins are represented by orange and violet bars respectively. The links represent the overlap between both DEGs and DEPs. The dashed lines indicate overlapped genes with arc correspondent colors. (b) miRNAs predicted to regulate the genes up-regulated in the transcriptomic dataset but not up-regulated in proteomics dataset. (c) Pathways enriched in transcriptomics and proteomics datasets. Numbers in red and blue are pathways enriched with up- and down-regulated genes respectively. Dashed lines indicate common pathways.

When considering the proteins that were exclusively found either in brain samples with CZS or in the ZIKV negative control brain, 99 shared up- and down-regulated genes and proteins were observed (Tables S3 and S4). These included genes such as LOX, PSMF1, NCAN, TNR, and NRCAM, which are associated with crosslinking of collagen and elastin, processing of class I MHC peptides, modulation of cell adhesion and migration, and neuronal cell adhesion.

We also integrated transcriptomics and proteomics at the pathway level. Gene Set Enrichment Analysis (GSEA) was performed using the mean foldchange between CZS and control brains as ranks and the Reactome pathways as gene sets. It was observed that a higher overlap between transcriptomics and proteomics with 47 pathways significantly enriched for both layers of information (Figure 3c). Furthermore, down-regulated pathways were again related to extracellular matrix organization and collagen formation and point to the central role of collagen in CZS outcome.

### Genetic variants associated with CZS

Whole exome sequencing analysis identified several rare variants with potential deleterious functions in five neonates (Z01, Z02, Z04, Z06, and Z07 in Figure 1). Combining variants that are presented in the same gene, it was found that 23 genes have at least one single nucleotide polymorphisms (SNP) in all five neonates (Figure 4a and Table S5). Variants in genes associated with extracellular matrix organization (collagen genes, *FBN2, FBN3*, and *FN1*) (Figure 4b and Table S5), as well as CNS development (*PTPRZ1*), immune system (*C7, C8A, IL4R, IL7, IRF3*, and *TLR2*), muscular contraction and arthrogryposis (*PIEZO2, RYR1* and *TTN*), and Notch and Wnt signaling pathways (*NOTCH3, NOTCH4* and *VANGL1*) were also found (Table S5).

**Figure 4.**
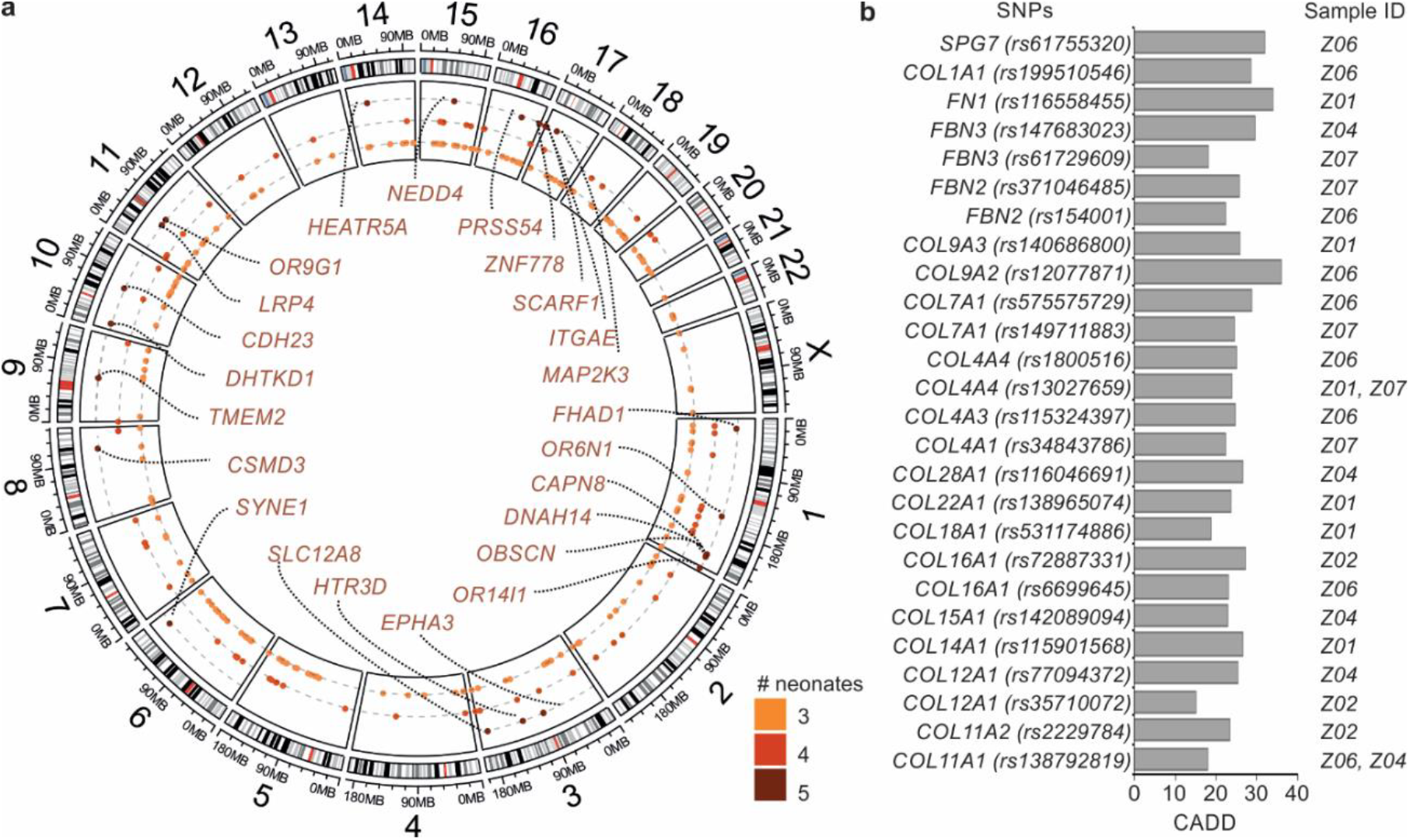
Single nucleotide polymorphisms in neonates with CZS. The exome analysis of five CZS cases (a) Genomic map showing genes with SNPs (MAF < 0.05 and CADD > 15) in three or more neonates with CZS. The outermost layer represents the reference genome (GRCh38). In the middle layer, each row represents genes with at least one SNP in three to five neonates. Dark brown rows represent genes that contain variants in all five neonates. (b) Most deleterious SNPs found in extracellular matrix genes.

### Integration of 3 Omics data types

The three layers of biological information were ultimately integrated into a network containing the gene variants and the RNAs and proteins differentially expressed in CZS cases (Figure 5). Only three genes appeared associated with CZS in all the layers—*COL1A1, COL12A1*, and *PTPRZ1* (Figure 5a). The former two collagen genes are related to the extracellular matrix organization. The latter gene *PTPRZ1* is instrumental to the differentiation of oligodendrocytes^22^ and have been associated with schizophrenia^23^. In total, there were 1,628 genes associated with CZS at either the genomic, transcriptomic, or proteomic level. Protein-protein interaction (PPI) data was used to construct a network with 341 of these genes (Figure 5b). Network analysis revealed several modules associated with proteasome degradation, axon guidance, the FGF signaling pathway, and Parkinson’s disease (Figure 5b).

**Figure 5.**
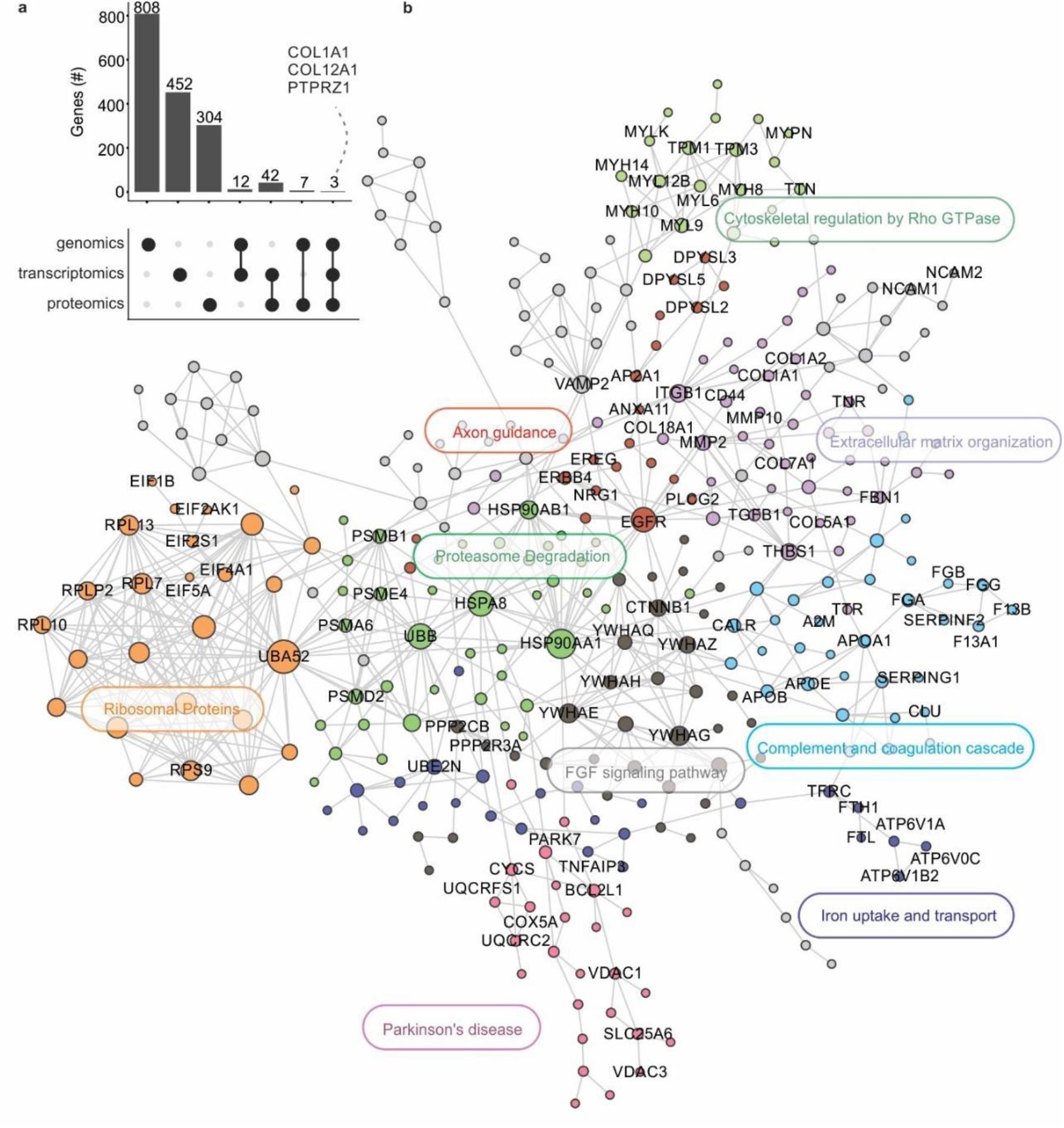
Integration of three molecular layers in *post-mortem* ZIKV-infected samples infected. (a) Intersections between three layers of information (genomics, transcriptomics, and proteomics) involved with CZS. Point diagram at the bottom represents the intersections between layers. Bar plot shows the number of genes in each intersection. A dashed line indicates the genes present in all the layers. (b) Protein-protein interaction network of CZS-related genes and their cellular pathways.

A more stringent analysis was performed considering only the 64 genes that were identified associated with CZS in at least two omics analyses (Figure 5a). Subsequently, these genes were integrated into a PPI network (Figure S2). Several central genes were observed—THBS1 promotes synaptogenesis^24^; DCN regulates collagen fibrils and matrix assembly^25^, and CLU shifts blood-brain barrier amyloid-beta drainage^26^.

Since all the analyses indicate that collagen genes are down-regulated in CZS brains, we performed a Gomori’s trichrome staining for total collagen in CZS brain, as well as in a different set of Zika negative control brains. We observed a reduction of collagen fibers in the CZS brains particularly in the adventitia of the vessels compared to Zika negative controls at the same gestational age. This reduction validated our transcriptome and proteome findings (Figure 6a). Next, the presence of COL1A1 was investigated through immunostaining directly in the brain tissues from CZS cases relative to the controls, which also showed less COL1A1 in all the CZS cases (Figure 6b). This corroborated the role of collagen isoforms in the neuropathogenesis associated with ZIKV infection in the brain tissues (Figure 6b).

**Figure 6.**
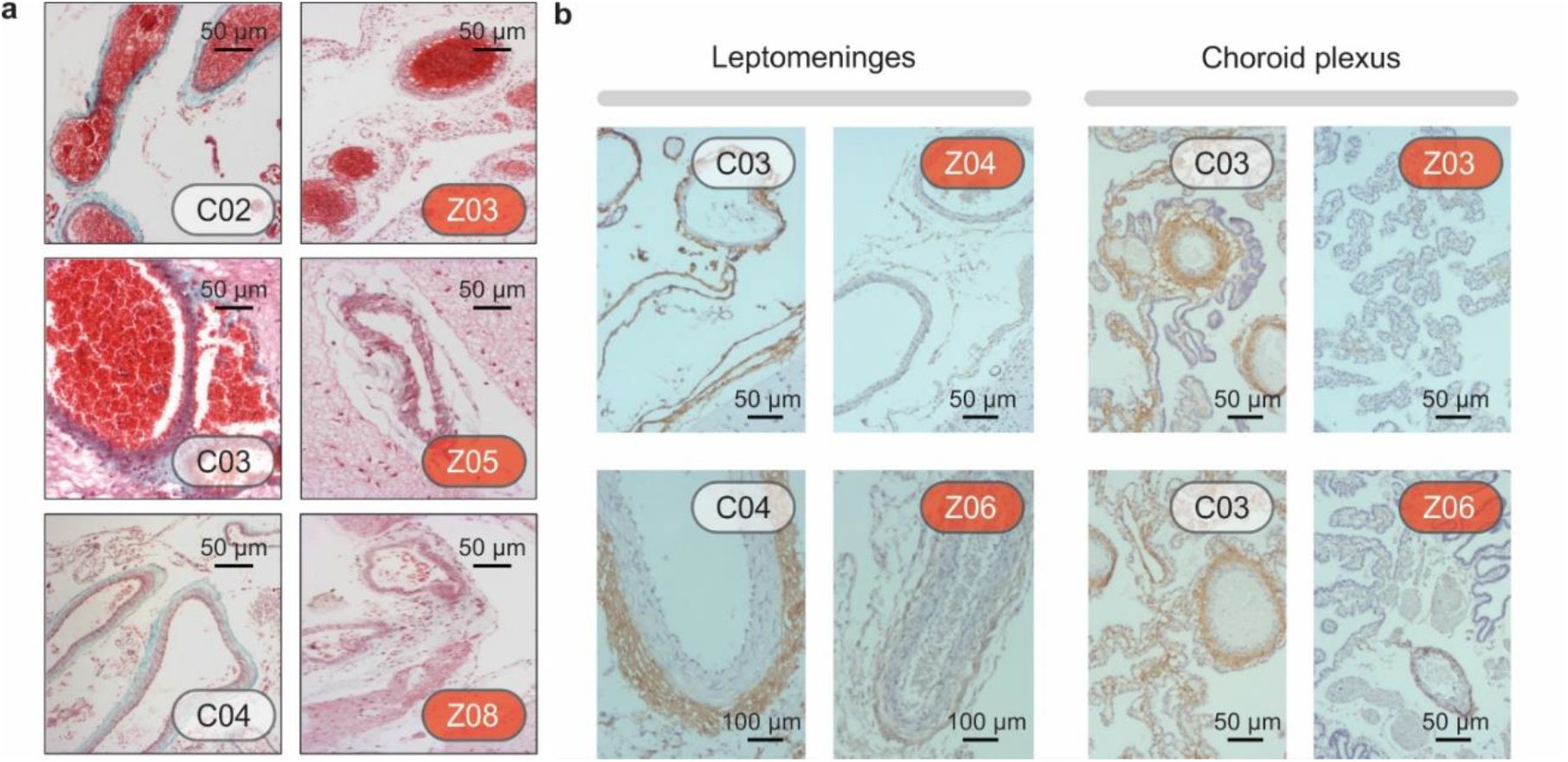
Reduction of collagen fibers in CZS cases compared with Zika negative controls. (a) Histopathological analysis confirms that the CZS brains have fewer collagen fibers compared to negative ZIKV control brains at the same gestational age. The total collagen that stains in green with the Gomori’s Trichrome is less evident than in controls, particularly in the adventitia of the vessels. (b) Immuno-histochemistry for collagen 1 also shows less collagen in CZS brains. Controls and CSZ cases are depicted as tables S1 and S3.

## Discussion

Our findings indicate that collagen genes and the extracellular matrix could play a significant role in CZS. Reduced levels of fibronectin and collagen IV increase the permeability of the blood-brain barrier^27^. Once this barrier is transposed, ZIKV could reach developing neural progenitor cells and severely disrupt the neural development. However, a more direct effect on fibroblast cells in the surrounding vasculature^28^ could not be discarded, and this effect could be a result of cell death or dysregulation of ECM expression or tissue deposition. The unique description of ECM gene modulation and ZIKV were reported in monkey experimental model during CNS viral persistence. Aid *et al.* showed that viral loads and viral persistence were negatively correlated with ECM genes, including collagen family genes^29^. Experiments using animal models indicate that deficiency in collagen compromise vessel resistance^30^. Mutations in collagen IV and fibronectin have induced impaired basement membranes or mesoderm defects respectively^31^. Moreover, mutations on *COL1A* and *COL4A1* caused defects in the basal membrane, resulting in a weakening of the brain vessels, arterial rupture and ischemic stroke^32,33^. Along with collagen isoforms, the down-regulation of the *LOX* gene that is responsible for cross-linking collagen fibers to elastin could potentialize the vascularity deficiency. This could explain the blood congestion in leptomeninges observed in all the brain samples analyzed here. Specifically, glycine mutations affecting exon 49 of the *COL1A2* gene was associated with an increased risk of intracranial bleeding^34^. Both collagen and *LOX* genes are stimulated in glioblastoma cells, and the suppression of this pathway by ZIKV infection could explain the decreasing of angiogenesis and anti-cancer effects that several authors are exploring to treat glioblastoma^35^-^37^ with ZIKV-like particles.

Mutations in type I collagen also affect the extracellular matrix by decreasing the amount of secreted collagen(s) impairing molecular and supramolecular assembly through the secretion of mutant collagen or by inducing endoplasmic reticulum stress and the unfolded protein response^38^. Mutated *COL1A1* were also associated with osteogenesis imperfecta, a generalized disorder of connective tissues that resembles the observed arthrogryposis phenotype common to all cases included in this work^39^. Mutations in *COL1A1/2* genes were associated with congenital brittle bones with the development of microcephaly and cataracts, as observed in the most severe cases of CSZ^40^. A dominant mutation in *COL12A1* was also related to joint laxity^41^, a phenotype often found in ZIKV-infected children^42^.

Cell-cell interaction is necessary for neuron migration through cortex layers during neurodevelopment^43^. L1CAM family of cell adhesion molecules are associated with neurite outgrowth and axon guidance^44^. In ZIKV-infected brains, NCAM1 and NFASC were up-regulated both at the RNA and protein levels. In addition, we found a rare variant in Neuronal Cell Adhesion Molecule gene (*NRCAM*) that could corroborate with ZIKV-infected brains phenotypes. These findings indicate that those genes/proteins could be the molecular basis for neurons migration defects already described by our group^10^ and should lead to CNS structural defects and reduction of cortical region observed in CZS newborns.

Among the pathways enriched for the up-regulated genes in ZIKV-infected samples, we found genes related to glutamate neurotransmitter release cycle and unblocking of NMDA receptor, glutamate binding, and activation. Previous experimental work revealed that NMDA receptor blockage has a protective effect on ZIKV-induced cell death^17^. In addition to gene expression, it was found that the genes associated with apoptosis were also up-regulated at the protein level. This corroborates the increased cell death proposed to neural progenitor cell pool and revealed by experimental data^45^.

Successful viral infection and disease must overcome the organism immune response. Pleiotrophin (PTN) is a cytokine that modulates inflammation in the CNS^46^. Additionally, PTN negatively regulates protein tyrosine phosphatase zeta (PTPRZ1), which binds to developmental proteins such as beta-catenin^47^. The results of this study showed that PTPRZ1 was up-regulated in ZIKV-infected brains both in the RNA and protein levels. Impressively, this gene also presented rare polymorphisms associated that raises the possibility of PTN–PTPRZ1 regulatory dysregulation and genetically driven suppression of neuroinflammation, which might result in a viral evasion mechanism. Considering the exclusive proteins (Table S3), another gene found in all three omics layers was NRCAM. This gene is a cell adhesion molecule that can interact with PTPRZ1^48^.

We also observed rare mutations in genes related to the immune system, including *IRF3* and *IL4R*. IRF3 plays a critical role in the innate immune response against DNA and RNA viruses, driving the transcription of type-I IFN genes^49^. Additionally, a mutation in the *IRF3* gene was associated with increased susceptibility to HSV-1 infection in the CNS in humans^50^. SNPs in the interleukin-4 receptor (IL4R) were also associated with increased susceptibility to dengue^51^. Our findings indicate that mutations in those genes could also confer increased susceptibility to ZIKV infection and CZS.

Altogether, this work is the first to investigate the molecular basis of ZIKV infection after vertical transmission using post-mortem brain samples. Despite the small sample size, these brain samples are unique considering the decrease in CZS cases worldwide. Our systems biology approach allowed us to unveil the different layers of biological information associated with CZS.

## Acknowledgments

We would like to thank Diego Santos and Heliomar Pereira Marcos who performed the collagen staining and immunohistochemistry. R.S.A. and A.M. are grateful to Biometrix and Dia.Pro Diagnostic Bioprobes, for the donation of ELISA kits for this project.

## Author contributions

EV and RDM performed the proteomics experiments, FCSN and GBD developed the proteomics rational, did data search, physiological analysis. FP, IJA, MYNJ, PLH, ASS, VS, HIN performed and analyzed the transcriptomics experiments. GLM, JBC, ALG, LWPA performed and analyzed the exome experiments. LC performed the neuropathological analysis (collagen staining and immuno-histochemistry). RSA, FP, ZFMV, GBD, ATRV, HIN integrated the omics datasets, interpreted the results and wrote the initial draft. All authors helped with the writing of the manuscript.

## Supplementary Material and methods

### Patients and neuroimaging studies

From June 2015 to July 2016, pregnant women presenting acute febrile illness clinic with a rash, fetal central nervous system (CNS) abnormalities at prenatal ultrasonography (US), and/ or postnatal microcephaly or other CNS malformation that was believed to be characteristic of congenital infection were referred to the Microcephaly Reference Center IPESQ in Campina Grande (Paraíba, Brazil) or Instituto Fernandes Figueira – Fiocruz (Rio de Janeiro, Brazil). This study includes imaging and autopsy data from an institutional review board–approved study (52888616.4.0000.5693 and 52675616.0.000.5269) that allowed for imaging and follow-up of presumed Zika virus infection in pregnant women and their neonates. Written informed consent was obtained from the pregnant women and/or the parents of neonates. Detailed demographic, medical, and prenatal history information, as well as clinical findings, were entered into case-report forms by multidisciplinary medical teams. All women were referred for at least one fetal ultrasonography during gestation. The onset symptoms included fever, exanthema, arthralgia, conjunctivitis, and headache in the pregnant women during gestation. The CNS of eight neonates who died in the first 48 h of life (two of them immediately after delivery), three from northeastern (Campina Grande, Paraíba state) and five from southeastern of Brazil (Rio de Janeiro) whose mothers reported typical symptoms of ZIKV infection until the 18th gestational week, were examined postmortem. Intrauterine fetal development was followed with ultrasonography and fetal MRI. Just after birth, the cephalic perimeter was measured and the percentile was calculated according to the expected for the gestational age 1. Prenatal US was performed by fetal medicine specialists using either a Voluson E8 unit (General Electric, Milwaukee, Wis) with transvaginal probes or a Samsung XG or WS80 unit (Samsung, Seoul, South Korea) with 2–9-MHz probes. MR imaging of the fetus was performed with a 3-T Skyra unit (Siemens Healthcare, Erlangen, Germany) or a 1.5-T Espree unit (Siemens Healthcare) with an eight-channel body coil and standard acquisition protocols. Postnatal head CT was performed with a 16-section CT scanner (Siemens Healthcare). Postnatal MR imaging was performed with a 1.5-T Espree brain MR imaging unit (Siemens Healthcare). Brain tissue images were acquired with a 64-channel multisection CT scanner (GE Healthcare) and a 3-T MR imaging unit (Achieva; Philips, Best, the Netherlands).

### Autopsies

Full autopsies were performed and the brains were fixed in 10% buffered formalin. In the three cases from IPESQ, one hemisphere was stored in RNA later and then frozen for virus RNA detection and transcriptome/proteomic analysis. In seven cases, the whole spinal cords were also removed, four of them with dorsal nerve ganglia (DRG). The upper cervical spinal cord was also sampled in two other cases, one with DRG. Formalin-fixed brains were weighed and the percentile was calculated according to the expected for the gestational age ^1^. In addition, samples from skeletal muscle (paravertebral, psoas, diaphragm or adjacent to the head of the femur) were taken and examined histologically in five cases. After macroscopic examination, representative areas, including those with macroscopic lesions, were processed for paraffin embedding and 5 μm histological sections were stained with hematoxylin and eosin (H&E). The neuropathological findings of these patients have been reported previously ^2^. Brains of ZIKV, CHIKV, DENV or STORCH negative controls of the same gestational age (30-41st) were obtained from Maternidade Escola - UFRJ (Rio de Janeiro, Brazil) covered by the institutional review board-approved study (1705093) and from Paraiba state. The death cause of correlate negative controls cases was genetic (trisomy of chromosome 18), acute perinatal anoxia, or complications of prematurity.

### Zika virus diagnostic procedures

ZIKV RNA was investigated in the mothers or babies through RT-PCR targeting the env gene as described by Lanciotti et al., 2008 ^3^. ZIKV RNA was detected in fluid samples including blood, urine, amniotic fluid obtained by amniocentesis during gestation, or in other fluids after birth (amniotic fluid and/or blood cord). ZIKV virus genome was also investigated postnatal in the autopsied tissues (placenta, brain, and other organs). Viral RNA was extracted from 140 μl fluids using QIAmp MiniElute Virus Spin (QIAgen, Hilden, Germany), following the manufacturer’s recommendations. ZIKV RNA detection was performed using One Step TaqMan RT-PCR (Thermo Fisher Scientific, Waltham, MA, United States) on 7500 Real-time PCR System (Applied Biosystems, Foster City, CA, United States) with primers, probes, and conditions as described elsewhere [4]. Fifty milligrams of frozen organs such as cerebral cortex, heart, skin, spleen, thymus, liver, kidneys, lung, and placenta were disrupted using Tissuerupter ® (QIAgen, Hilden, Germany) using 325 μl of RTL buffer from RNEasy Plus Mini Kit (QIAgen, Hilden, Germany), following the manufacturer’s protocol. RNA extraction was processed with Rneasy Plus Mini Kit (QiAgen, Hilden, Germany), following the recommendations of the manufacturer. Real-time RT-PCR was performed using 1 μg of total tissue RNA using One Step TaqMan RT-PCR (Thermo Fisher Scientific, Waltham, MA, United States) as described above. Dengue and Chikungunya virus infections were excluded in all cases (fluids and tissues) either by RT-PCR using ZDC Trioplex kits (Bio-Manguinhos, Fiocruz, Rio de Janeiro, Brazil) or serological ELISA for qualitative determination of IgM and IgG (Kit XGen, Biometrix, Brazil and Euroimmum kit, Lübeck, Germany). Other congenital pathogens including Syphilis, Cytomegalovirus, Herpes Virus 1/2, Toxoplasma Gondii and Rubella Virus (STORCH) were discharged by serological ELISA against IgM (Dia.Pro Diagnostic Bioprobes, Italy), following the manufacturer’s recommendations.

ZIKV RNA in situ hybridization (ISH) was also investigated on formalin-fixed paraffin embedded (FFPE) tissue sections of all brain tissues using two commercial RNAscope Target Probes (Advanced Cell Diagnostics, Hayward, CA, United States) catalog # 464531 and 463781 complementary to sequences 866-1763 and 1550-2456 of ZIKV genome, respectively. Pretreatment, hybridization, and detection techniques were performed according to the manufacturer’s protocols ^2^.

### Collagen staining/Immunohistochemistry

For total collagen visualization, paraffin-embedded sections from formalin fixed fragments of post-mortem brains were stained with the Gomori’s Trichrome reagent.

From the leptomeninges, choroid plexus of ZIKV cases and controls immuno-histochemical reactions were performed, using the following monoclonal antibody (Sigma) and dilution: anti-collagen type 1, clone col-1, 1,1000. Briefly, 5 μm thick tissue sections were incubated in an oven at 37 °C for six hours, deparaffinized in xylene and rehydrated by placing in decreasing concentrations of alcohol and washed in distilled water. To enhance antigen retrieval, the tissue sections were pretreated in a pressure cooker for 15 minutes in the solution 1/20 Declare (pH 6) / 1/100 Trilogy (pH9) in distilled water. To block endogenous peroxidase activity, they were exposed to hydrogen peroxide, washed with distilled water and rinsed in phosphate buffered saline (PBS) to stop enzymatic digestion, then incubated with the primary antibody overnight at 4°C, rinsed in PBS for 5 minutes and incubated with Polymer Hi-Def (horseradish peroxidase system) for 10 minutes at room temperature preceded by several washes in PBS. The peroxidase reaction was visualized with DAB substrate, rinsed in running water; the sections were then counterstained with Meyer’s hematoxylin for 1 minute, washed in running tap water for 3 minutes, dehydrated in alcohol, cleared in xylene and mounted in a resinous medium.

### Library preparation and RNA-sequencing

Brain samples were frozen in RNALater (Ambion®) and stored at −80 ° C until extraction. The tissue was broken and homogenized by TissueRupter® (QIAGEN) and RNA extraction was performed with the RNAeasy Plus Mini® kit (QIAGEN), following the protocol suggested by the manufacturer.

The integrity of RNA was evaluated using Agilent 2100 Bioanalyzer with RNA 6000 Pico. Total RNA was quantified by Quant-iT ™ RiboGreen® RNA reagent and Kit (Invitrogen, Life Technologies Corp.) and the cDNA library was constructed following the SMARTer Stranded Total RNASeq Peak Input Mammalian Kit protocol (Takara Bio USA). The size distribution of the cDNA library was measured by 2100 Bioanalyzer and quantitated prior to sequencing using Quant-iT™ PicoGreen® RNA reagent and Kit (Invitrogen, Life Technologies Corp.). The libraries were diluted to 4nM with 15% PhiX. The cDNA library was sequenced with MySeq System (Illumina®, San Diego, CA) using the MiSeq Reagent kit (150 cycles, 2 x 75 paired-end).

### Pre-processing and analysis of RNA-seq data

FASTQ quality control was performed using FastQC tool (https://www.bioinformatics.babraham.ac.uk/projects/fastqc/). Paired-end reads were aligned to the human genome, ENSEMBL GRCh38.89, by STAR v.2.5.3.a, an ultra-fast aligner ^4^. Then, aligned reads were quantified using featureCounts v.1.5.3 ^5^. Differentially expressed genes (DEGs) between control and infected conditions were detected using DESeq2 v.1.16.1 R package ^6^ (adjusted p-value < 0.1). Functional enrichment analysis was performed using the Reactome pathways (https://reactome.org/) and the EnrichR tool^7^ for Over Representation Analysis. The overlap between gene lists was performed using circlize ^8^ and UpSetR ^9^ packages.

### DNA extraction for Whole Exome Sequencing

DNA extraction and exome sequencing Genomic DNA was extracted from the central nervous system. Exome sequencing libraries were prepared using Illumina TruSeq® Exome Kit (8 rxn × 6plex). Sequencing was performed using Illumina NextSeq® 500/550 High Output Kit v2 (150 cycles), generating 2×75 bp paired-end reads.

### Whole Exome Sequencing analysis

The quality of the exome libraries was evaluated using the FASTQC tool (http://www.bioinformatics.babraham.ac.uk/projects/fastqc/). The removal of reads or fragments with low quality was performed by the Trimmomatic software (http://www.usadellab.org/cms/?page=trimmomatic). The resulting high-quality reads were aligned with the human genome as a reference (version GRCh38) using Bowtie2 ^10^ with very sensitive default preset (-D 20 -R 3 -N 1 -L 20 -i S,1,0.50), except to one mismatch per seed region (-N 1). The optical duplicates were marked with mark duplicates tool (http://broadinstitute.github.io/picard/). Further, the Genome Analysis Toolkit (GATK) version 3.7 ^11^ was used to call Single Nucleotide Variants (SNVs), small insertions and deletions (INDELs). All variants were annotated with the HaplotypeCaller following the GATK best practices manual ^12,13^. The variant calls with a read coverage of ≤ 5 reads or a MAP quality (MAPQ) of ≤ 30 were filtered out in order to avoid false positives. The SnpEff ^14^ and SnpSift ^15^ version 4.3r tools were used to predict and annotate the functional impact of variants, using the dbSNP (build 151) ^16^ and dbNSFP (version 3.5) ^17^ databases. The variants with MAF (minor allele frequency) ≤ 5% in at least one of the following databases (retrieved from dbNSFP database V3.5) were considered: 1000 Genomes project Phase 3 (http://www.internationalgenome.org/), ExAC (http://exac.broadinstitute.org/), gnomAD(http://gnomad.broadinstitute.org/), TOPMed (https://www.nhlbiwgs.org/), ESP6500 (http://evs.gs.washington.edu/EVS/), TwinsUK (http://www.twinsuk.ac.uk/), ALSPAC (http://www.bris.ac.uk/alspac/), ABraOM (http://abraom.ib.usp.br/). We also verified the presence of variants in GWAS catalog retrieved from https://www.ebi.ac.uk/gwas/ and CLINVAR - release 20180603 (https://www.ncbi.nlm.nih.gov/clinvar/). The 1000 Genomes project Phase 3, EXAC and gnomAD included African, Ad Mixed American, East Asian, Europea, South Asian and Non-Finnish European populations. The TOPMed and ESP6500 included cohorts from the United States. The TwinsUK included old aged twins from the United Kingdom and ALSPAC included European cohorts. The ABraOM is a variant repository comprising a cohort of elderly Brazilians ^18^. We considered only the variants with CADD score ≥ 15 ^19^ and used a set of functional effect predictors such as MetaSVM, FATHMM, LRT, PROVEAN, Polyphen2-HDIV, Polyphen2-HVAR, MutationTaster, Mutation Assessor and SIFT for variants prioritization ^20^. All variants of interest were manually inspected with IGV tool ^21^.

### Protein extraction

Approximately thirty milligrams of brain tissue was homogenized with 1.5 ml of extraction solution containing 5% of sodium deoxycholate (SDC), 0.75 mM dithiothreitol (DTT), protease and phosphatase inhibitors (Roche) in a TissueRuptor (QIAgen). After incubation for 20 min at 80 °C, the solution was vortexed for 20 s and centrifuged for 30 min at 4 °C, 20,000 g. The pellet from overnight precipitation of 400 μl of the supernatant with cold acetone (ratio 1: 4), was washed two times with acetone, centrifuged for 15 min at 4 ° C, 20,000 *g* and dried. After solubilizing with 7M urea / 2M thiourea with 2 % SDC we used the Qubit® protein assay kit (Invitrogen) to measure protein content according to the manufacturer’s instructions.

### Enzymatic digestion

Reduction and alkylation of 100 μg of soluble proteins used 10 mM DTT for 1 h at 30 °C and 40 mM IAA for 30 min at room temperature, in the dark. Samples were diluted 1:10 with triethylammonium bicarbonate buffer (TEAB) 100 mM pH 8.5 and digested with trypsin (1:25, w/w) for 18 hours at 35 °C. Addition of a final concentration of 1% TFA stopped digestion and two centrifugations for 15 min, 4 °C at 20,000 g removed SDC. Finally, samples were desalted in Macro SpinColumns C_18_ (*Harvard Apparatus*) and dried in a vacuum concentrator (Martin Christ). Peptides were suspended in 15 μl of formic acid 0.1% and quantified by the Qubit ® protein assay as described by the manufacturer.

### Nano-LC MS^2^ analysis

Each sample was analyzed four times (4 technical replicates) in an EASY 1000 - nLC (Thermo Scientific) coupled to a Q-Exactive Plus mass spectrometer (Thermo Scientific). Two µg of the peptide mixture was loaded in a homemade 3 cm trap column, 200 µm I.D., 5 µm ReprosilPur C18 AQ (Dr. Maishy) beads and fractionated in 20 cm Self-Pack PicoFrit analytical column (New Objective), 75 µm I.D., 3 µm ReprosilPur C18 AQ (Dr. Maishy). nLC gradient fractionation lasted 180 min and a flow-rate of 250 nL/min: 167 min from 5% to 40% of solvent B (95% ACN/ 5% H_2_O / 0.1% formic acid); 5 min from 40% to 95% of solvent B; and 8 min in 95% of solvent B. Column and trap were equilibrated with solvent A (95% H_2_O / 5% ACN / 0.1% formic acid) after each run for 15 and 2 min, respectively. The instrument was set in the positive polarity and Full-MS/DD MS^2^ mode. Selected full scan parameters were 1 microscan, 70,000 resolution at 200 *m/z*, 3e^6^ ions for AGC target, 50 ms maximum injection time and range of 375-2000 *m/z*. Top 20 DD-MS2 parameters were 17,000 resolution, 200 *m/z*, 1e^5^ ions for AGC target, the maximum injection time of 100 ms, 1.2 Th of isolation window, NCE of 30, minimum intensity threshold of 10,000 ions, and dynamic exclusion of 60 s.

### Proteomics analysis

For database search, raw data were processed using Proteome Discoverer 2.1 (PD2.1) software (Thermo Scientific) and the SuperQuant strategy performed by nodes MSn-Deconvolution and Complementary Finder as referred in ^22^. Search performed against all reviewed human and virus entries present in the UniProt Database (Jan/2017) used Sequest HT algorithm. Virus proteins were not considered for the analyses. The parameters used for the search were full tryptic peptides, two missed cleavages allowed, precursor mass tolerance of 10 ppm, 0.1 Da product ion mass tolerance, cysteine carbamidomethylation as fixed modification, and methionine oxidation and protein N-terminal acetylation as variable modifications. To estimate the False Discovery Rate (FDR) of <1% we used the node Percolator present in the PD2.1 using maximum parsimony. A cutoff score was established to accept a false-discovery rate (FDR) of 1% at the protein and peptide level, and proteins were grouped in master proteins using the maximum parsimony principle.

Quantification used the workflow node Precursor Ions Area Detector in PD2.1. The peak area estimated by the Extracted Ion Chromatogram (XIC) for the three most abundant distinct peptides of each protein were averaged and used for relative quantification. Statistical analysis was carried out on Perseus version 1.6.0.7. ^23^. Data was converted to log2 scale and normalized by subtracting the converted protein area value (XIC) by the median of the sample distribution. Only proteins with peak area averages present in at least three runs were used for quantitative evaluation.

We used the limma R package ^24^ to identify the proteins that were up- or down-regulated in CZS brains compared to the control brain. A cutoff Adjusted P-value < 0.1 was used. Proteins detected in at least 2 CZS samples and not detected in the control were considered exclusively expressed in CZS. Proteins detected in the control and not detected in any of the CZS samples were considered exclusively expressed in control. Functional enrichment analysis was performed using the Reactome pathways (https://reactome.org/) and the EnrichR tool^7^ for Over Representation Analysis.

### Network analysis

Protein-protein interaction (PPI) networks and the miRNA-gene network were generated using the NetworkAnalyst tool ^25^. Protein-protein interactions (edges) were retrieved from STRING interactome with confidence score 900. The miRNA-gene interaction data were collected from TarBase and miRTarBase (validated interactions). We used the Minimum Network tool to include the seed genes/proteins (i.e. DEGs or DEPs) as well as the essential non-seed genes/proteins that keep the network connection. Cytoscape program ^26^ was also used to visualize the networks.

---

**Figure S1.**
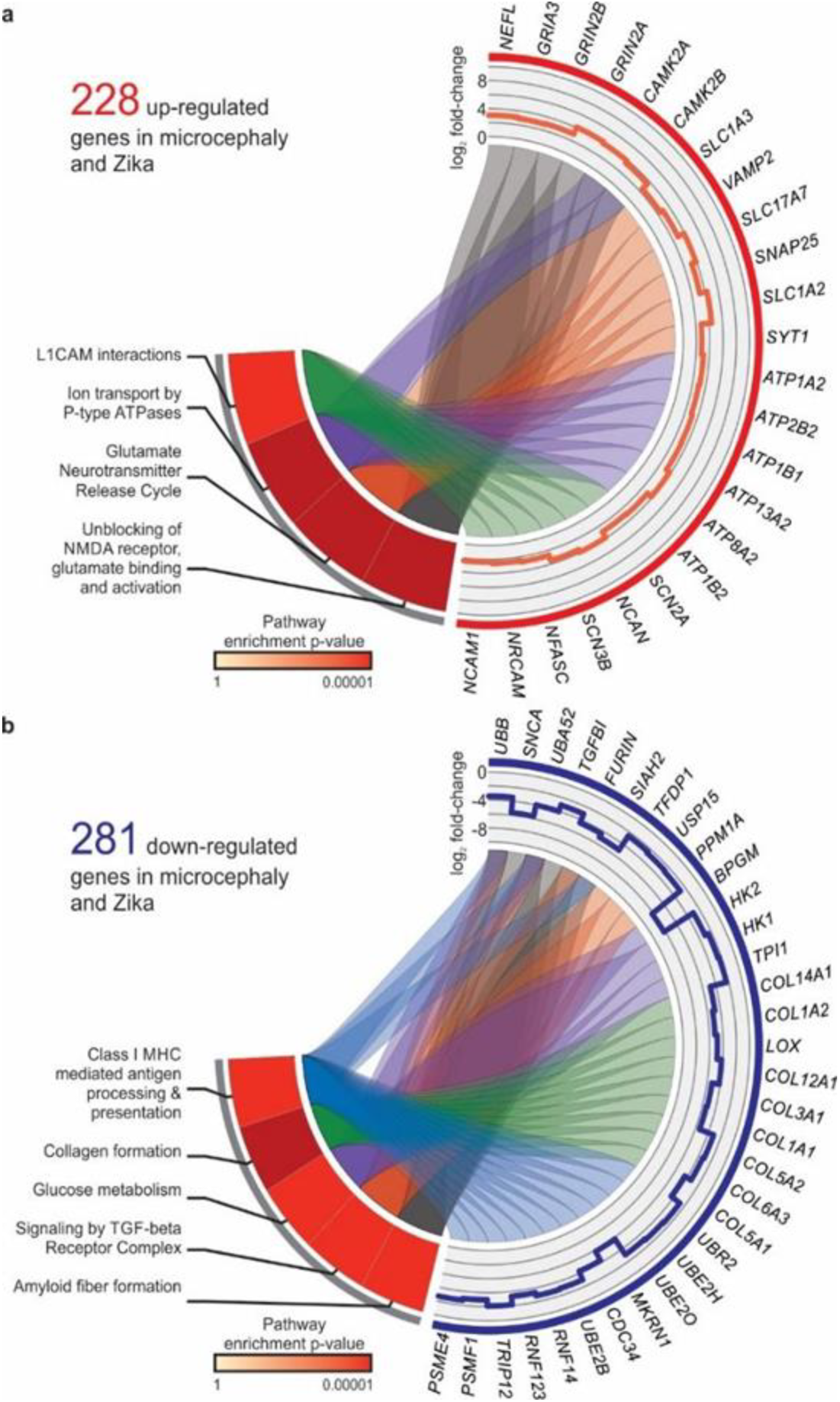
Modulation of brain-expressed genes in neonates with CZS evidenced by transcriptome. Genes and pathways up-regulated (a) or down-regulated (b) in CZS compared to ZIKV negative control brain in the prefrontal cortex. The pathways enriched by Over Representation Analysis (left) are present in the outermost layer and the differentially expressed genes (right) found in these pathways (up-regulated in red and down-regulated in blue). In the innermost layer, the links indicate the pathway in which the genes were found. In the middle layer, the colors in the heat map represent the pathway enrichment P-value obtained by Over Representation Analysis, while the line graph represents the log_2_ fold change value for each gene in the CZS samples relative to the control.

**Figure S2.**
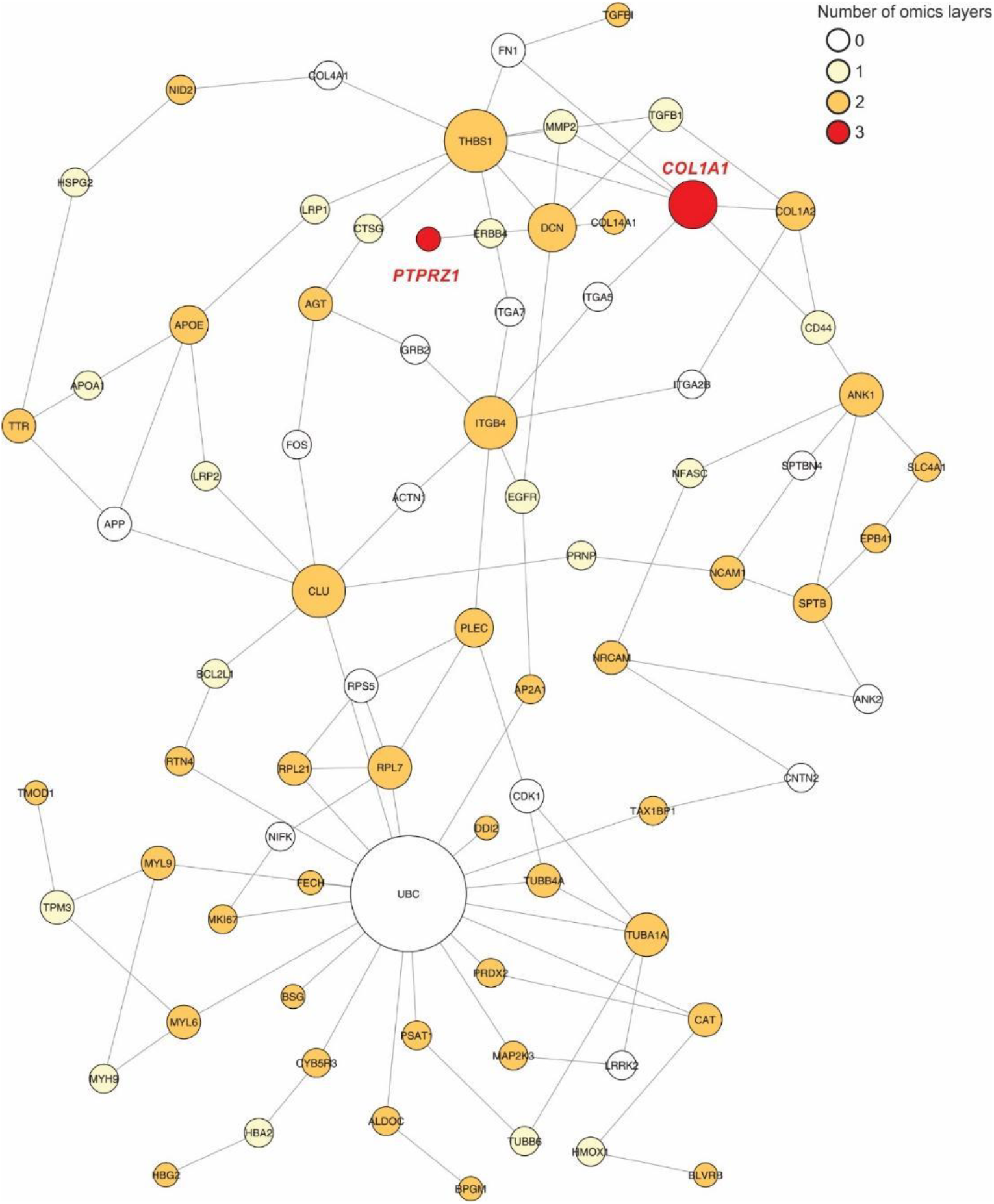
A network of highly associated CZS-related genes. More stringent criteria (at least two layers of biological information) was used to select the genes to construct the protein-protein interaction network.

## References

1 Melo, A. S. D. et al. Congenital Zika Virus Infection Beyond Neonatal Microcephaly. Jama Neurol 73, 1407–1416, doi:10.1001/jamaneurol.2016.3720 (2016).

2 Mlakar, J. et al. Zika Virus Associated with Microcephaly. New Engl J Med 374, 951–958, doi:10.1056/NEJMoa1600651 (2016).

3 Ventura, C. V., Maia, M., Bravo-Filho, V., Gois, A. L. & Belfort, R. Zika virus in Brazil and macular atrophy in a child with microcephaly. Lancet 387, 228–228, doi:10.1016/S0140-6736(16)00006-4 (2016).

4 Brasil, P. et al. Zika Virus Infection in Pregnant Women in Rio de Janeiro. N Engl J Med 375, 2321–2334, doi:10.1056/NEJMoa1602412 (2016).

5 Duffy, M. R. et al. Zika virus outbreak on Yap Island, Federated States of Micronesia. N Engl J Med 360, 2536–2543, doi:10.1056/NEJMoa0805715 (2009).

6 Johansson, M. A., Mier-y-Teran-Romero, L., Reefhuis, J., Gilboa, S. M. & Hills, S. L. Zika and the Risk of Microcephaly. N Engl J Med 375, 1–4, doi:10.1056/NEJMp1605367 (2016).

7 Moore, C. A. et al. Characterizing the Pattern of Anomalies in Congenital Zika Syndrome for Pediatric Clinicians. JAMA Pediatr 171, 288–295, doi:10.1001/jamapediatrics.2016.3982 (2017).

8 Tang, H. L. et al. Zika Virus Infects Human Cortical Neural Progenitors and Attenuates Their Growth. Cell Stem Cell 18, 587–590, doi:10.1016/j.stem.2016.02.016 (2016).

9 Shao, Q. et al. Zika virus infection disrupts neurovascular development and results in postnatal microcephaly with brain damage. Development 143, 4127–4136, doi:10.1242/dev.143768 (2016).

10 Chimelli, L. et al. The spectrum of neuropathological changes associated with congenital Zika virus infection. Acta Neuropathol 133, 983–999, doi:10.1007/s00401-017-1699-5 (2017).

11 Garcez, P. P. et al. Zika virus disrupts molecular fingerprinting of human neurospheres. Sci Rep 7, 40780, doi:10.1038/srep40780 (2017).

12 Kwissa, M. et al. Dengue virus infection induces expansion of a CD14(+)CD16(+) monocyte population that stimulates plasmablast differentiation. Cell Host Microbe 16, 115–127, doi:10.1016/j.chom.2014.06.001 (2014).

13 Nakaya, H. I. & Pulendran, B. Vaccinology in the era of high-throughput biology. Philos Trans R Soc Lond B Biol Sci 370, doi:10.1098/rstb.2014.0146 (2015).

14 Avey, S. et al. Multicohort analysis reveals baseline transcriptional predictors of influenza vaccination responses. Sci Immunol 2, doi:UNSP eaa14656 10.1126/sciimmunol.aal4656 (2017).

15 Hoen, B. et al. Pregnancy Outcomes after ZIKV Infection in French Territories in the Americas. N Engl J Med 378, 985–994, doi:10.1056/NEJMoa1709481 (2018).

16 Krow-Lucal, E. R. et al. Association and birth prevalence of microcephaly attributable to Zika virus infection among infants in Paraiba, Brazil, in 2015-16: a case-control study. Lancet Child Adolesc 2, 204–213, doi:10.1016/S2352-4642(18)30020-8 (2018).

17 Costa, V. V. et al. N-Methyl-d-Aspartate (NMDA) Receptor Blockade Prevents Neuronal Death Induced by Zika Virus Infection. MBio 8, doi:10.1128/mBio.00350-17 (2017).

18 Obermeier, B., Daneman, R. & Ransohoff, R. M. Development, maintenance and disruption of the blood-brain barrier. Nat Med 19, 1584–1596, doi:10.1038/nm.3407 (2013).

19 Mala, J. G. & Rose, C. Interactions of heat shock protein 47 with collagen and the stress response: an unconventional chaperone model? *Life Sci* 87, 579–586, doi:10.1016/j.lfs.2010.09.024 (2010).

20 Deckx, S., Heymans, S. & Papageorgiou, A. P. The diverse functions of osteoglycin: a deceitful dwarf, or a master regulator of disease? FASEB J 30, 2651–2661, doi:10.1096/fj.201500096R (2016).

21 Kozak, R. A. et al. MicroRNA and mRNA Dysregulation in Astrocytes Infected with Zika Virus. Viruses 9, doi:10.3390/v9100297 (2017).

22 Kuboyama, K. et al. Protein tyrosine phosphatase receptor type z negatively regulates oligodendrocyte differentiation and myelination. PLoS One 7, e48797, doi:10.1371/journal.pone.0048797 (2012).

23 Buxbaum, J. D. et al. Molecular dissection of NRG1-ERBB4 signaling implicates PTPRZ1 as a potential schizophrenia susceptibility gene. Mol Psychiatr 13, 162–172, doi:10.1038/sj.mp.4001991 (2008).

24 Christopherson, K. S. et al. Thrombospondins are astrocyte-secreted proteins that promote CNS synaptogenesis. Cell 120, 421–433, doi:DOI 10.1016/j.cell.2004.12.020 (2005).

25 Robinson, K. A. et al. Decorin and biglycan are necessary for maintaining collagen fibril structure, fiber realignment, and mechanical properties of mature tendons. Matrix Biol 64, 81–93, doi:10.1016/j.matbio.2017.08.004 (2017).

26 Wojtas, A. M. et al. Loss of clusterin shifts amyloid deposition to the cerebrovasculature via disruption of perivascular drainage pathways. P Natl Acad Sci USA 114, E6962–E6971, doi:10.1073/pnas.1701137114 (2017).

27 Baeten, K. M. & Akassoglou, K. Extracellular matrix and matrix receptors in blood-brain barrier formation and stroke. Dev Neurobiol 71, 1018–1039, doi:10.1002/dneu.20954 (2011).

28 Hamel, R. et al. Biology of Zika Virus Infection in Human Skin Cells. J Virol 89, 8880–8896, doi:10.1128/JVI.00354-15 (2015).

29 Aid, M. et al. Zika Virus Persistence in the Central Nervous System and Lymph Nodes of Rhesus Monkeys. Cell 169, 610–620 e614, doi:10.1016/j.cell.2017.04.008 (2017).

30 Kuo, D. S., Labelle-Dumais, C. & Gould, D. B. COL4A1 and COL4A2 mutations and disease: insights into pathogenic mechanisms and potential therapeutic targets. Hum Mol Genet 21, R97–110, doi:10.1093/hmg/dds346 (2012).

31 Poschl, E. et al. Collagen IV is essential for basement membrane stability but dispensable for initiation of its assembly during early development. Development 131, 1619–1628, doi:10.1242/dev.01037 (2004).

32 Dittrich, R. et al. Connective tissue and vascular phenotype in patients with cervical artery dissection. Neurology 68, 2120–2124, doi:10.1212/01.wnl.0000264892.92538.a9 (2007).

33 Weng, Y. C. et al. COL4A1 mutations in patients with sporadic late-onset intracerebral hemorrhage. Ann Neurol 71, 470–477, doi:10.1002/ana.22682 (2012).

34 Faqeih, E., Roughley, P., Glorieux, F. H. & Rauch, F. Osteogenesis imperfecta type III with intracranial hemorrhage and brachydactyly associated with mutations in exon 49 of COL1A2. Am J Med Genet A 149A, 461–465, doi:10.1002/ajmg.a.32653 (2009).

35 Zhu, Z. et al. Zika virus has oncolytic activity against glioblastoma stem cells (vol 214, pg 3145, 2017). Journal of Experimental Medicine 214 (2017).

36 Chen, Q. et al. Treatment of Human Glioblastoma with a Live Attenuated Zika Virus Vaccine Candidate. MBio 9, doi:10.1128/mBio.01683-18 (2018).

37 Mammoto, T. et al. Role of collagen matrix in tumor angiogenesis and glioblastoma multiforme progression. Am J Pathol 183, 1293–1305, doi:10.1016/j.ajpath.2013.06.026 (2013).

38 Bateman, J. F., Boot-Handford, R. P. & Lamande, S. R. Genetic diseases of connective tissues: cellular and extracellular effects of ECM mutations. Nat Rev Genet 10, 173–183, doi:10.1038/nrg2520 (2009).

39 Marini, J. C. et al. Consortium for osteogenesis imperfecta mutations in the helical domain of type I collagen: regions rich in lethal mutations align with collagen binding sites for integrins and proteoglycans. Hum Mutat 28, 209–221, doi:10.1002/humu.20429 (2007).

40 Plotkin, H. Syndromes with congenital brittle bones. BMC Pediatr 4, 16, doi:10.1186/1471-2431-4-16 (2004).

41 Zou, Y. et al. Recessive and dominant mutations in COL12A1 cause a novel EDS/myopathy overlap syndrome in humans and mice. Hum Mol Genet 23, 2339–2352, doi:10.1093/hmg/ddt627 (2014).

42 Siqueira, W. L., Moffa, E. B., Mussi, M. C. M. & Machado, M. A. D. M. Zika virus infection spread through saliva - a truth or myth? Braz Oral Res 30, doi:ARTN e46 10.1590/1807-3107BOR-2016.vol30.0046 (2016).

43 Nadarajah, B., Alifragis, P., Wong, R. O. L. & Parnavelas, J. G. Neuronal migration in the developing cerebral cortex: Observations based on real-time imaging. Cereb Cortex 13, 607–611, doi:DOI 10.1093/cercor/13.6.607 (2003).

44 Maness, P. F. & Schachner, M. Neural recognition molecules of the immunoglobulin superfamily: signaling transducers of axon guidance and neuronal migration (vol 10, pg 19, 2007). Nat Neurosci 10, 263–263, doi:DOI 10.1038/nn0207-263b (2007).

45 Cugola, F. R. et al. The Brazilian Zika virus strain causes birth defects in experimental models. Nature 534, 267–271, doi:10.1038/nature18296 (2016).

46 Fernandez-Calle, R. et al. Pleiotrophin regulates microglia-mediated neuroinflammation. J Neuroinflamm 14, doi:ARTN 46 10.1186/s12974-017-0823-8 (2017).

47 Pariser, H., Ezquerra, L., Herradon, G., Perez-Pinera, P. & Deuel, T. F. Fyn is a downstream target of the pleiotrophin/receptor protein tyrosine phosphatase beta/xi-signaling pathway: Regulation of tyrosine phosphorylation of Fyn by pleiotrophin. Biochem Bioph Res Co 332, 664–669, doi:10.1016/j.bbrc.2005.05.007 (2005).

48 Sakurai, T. et al. Induction of neurite outgrowth through contactin and Nr-CAM by extracellular regions of glial receptor tyrosine phosphatase beta. J Cell Biol 136, 907–918, doi:DOI10.1083/jcb.136.4.907 (1997).

49 Liu, S. et al. Phosphorylation of innate immune adaptor proteins MAVS, STING, and TRIF induces IRF3 activation. Science 347, aaa2630, doi:10.1126/science.aaa2630 (2015).

50 Andersen, L. L. et al. Functional IRF3 deficiency in a patient with herpes simplex encephalitis. J Exp Med 212, 1371–1379, doi:10.1084/jem.20142274 (2015).

51 Useche, Y. M. et al. Association of IL4R-rs1805016 and IL6R-rs8192284 polymorphisms with clinical dengue in children from Colombian populations. J Infect Public Health, doi:10.1016/j.jiph.2018.08.009 (2018).

## References

1 Larroche, J.-C. Developmental pathology of the neonate. (Excerpta Medica ; sole distributors for the U.S.A. and Canada, Elsevier/North-Holland, 1977).

2 Chimelli, L. et al. The spectrum of neuropathological changes associated with congenital Zika virus infection. Acta Neuropathol 133, 983–999, doi:10.1007/s00401-017-1699-5 (2017).

3 Lanciotti, R. S. et al. Genetic and serologic properties of Zika virus associated with an epidemic, Yap State, Micronesia, 2007. Emerg Infect Dis 14, 1232–1239, doi:10.3201/eid1408.080287 (2008).

4 Dobin, A. et al. STAR: ultrafast universal RNA-seq aligner. Bioinformatics 29, 15–21, doi:10.1093/bioinformatics/bts635 (2013).

5 Shin, J. et al. Single-Cell RNA-Seq with Waterfall Reveals Molecular Cascades underlying Adult Neurogenesis. Cell Stem Cell 17, 360–372, doi:10.1016/j.stem.2015.07.013 (2015).

6 Love, M. I., Huber, W. & Anders, S. Moderated estimation of fold change and dispersion for RNA-seq data with DESeq2. Genome Biol 15, 550, doi:10.1186/s13059-014-0550-8 (2014).

7 Chen, E. Y. et al. Enrichr: interactive and collaborative HTML5 gene list enrichment analysis tool. BMC Bioinformatics 14, 128, doi:10.1186/1471-2105-14-128 (2013).

8 Gu, Z., Gu, L., Eils, R., Schlesner, M. & Brors, B. circlize Implements and enhances circular visualization in R. Bioinformatics 30, 2811–2812, doi:10.1093/bioinformatics/btu393 (2014).

9 Lex, A., Gehlenborg, N., Strobelt, H., Vuillemot, R. & Pfister, H. UpSet: Visualization of Intersecting Sets. IEEE Trans Vis Comput Graph 20, 1983–1992, doi:10.1109/TVCG.2014.2346248 (2014).

10 Langmead, B. & Salzberg, S. L. Fast gapped-read alignment with Bowtie 2. Nat Methods 9, 357–359, doi:10.1038/nmeth.1923 (2012).

11 McKenna, A. et al. The Genome Analysis Toolkit: a MapReduce framework for analyzing next-generation DNA sequencing data. Genome Res 20, 1297–1303, doi:10.1101/gr.107524.110 (2010).

12 DePristo, M. A. et al. A framework for variation discovery and genotyping using next-generation DNA sequencing data. Nat Genet 43, 491-+, doi:10.1038/ng.806 (2011).

13 Van der Auwera, G. A. et al. From FastQ data to high confidence variant calls: the Genome Analysis Toolkit best practices pipeline. Curr Protoc Bioinformatics 43, 11 10 11–33, doi:10.1002/0471250953.bi1110s43 (2013).

14 Cingolani, P. et al. A program for annotating and predicting the effects of single nucleotide polymorphisms, SnpEff: SNPs in the genome of Drosophila melanogaster strain w(1118); iso-2; iso-3. Fly 6, 80–92, doi:10.4161/fly.19695 (2012).

15 Cingolani, P. et al. Using Drosophila melanogaster as a Model for Genotoxic Chemical Mutational Studies with a New Program, SnpSift. Front Genet 3, 35, doi:10.3389/fgene.2012.00035 (2012).

16 Sherry, S. T. et al. dbSNP: the NCBI database of genetic variation. Nucleic Acids Res 29, 308–311 (2001).

17 Liu, X., Jian, X. & Boerwinkle, E. dbNSFP: a lightweight database of human nonsynonymous SNPs and their functional predictions. Hum Mutat 32, 894–899, doi:10.1002/humu.21517 (2011).

18 Naslavsky, M. S. et al. Exomic variants of an elderly cohort of Brazilians in the ABraOM database. Hum Mutat 38, 751–763, doi:10.1002/humu.23220 (2017).

19 van der Velde, K. J. et al. GAVIN: Gene-Aware Variant INterpretation for medical sequencing. Genome Biol 18, 6, doi:10.1186/s13059-016-1141-7 (2017).

20 Liu, X., Wu, C., Li, C. & Boerwinkle, E. dbNSFP v3.0: A One-Stop Database of Functional Predictions and Annotations for Human Nonsynonymous and Splice-Site SNVs. Hum Mutat 37, 235–241, doi:10.1002/humu.22932 (2016).

21 Robinson, J. T. et al. Integrative genomics viewer. Nat Biotechnol 29, 24–26, doi:10.1038/nbt.1754 (2011).

22 Gorshkov, V., Verano-Braga, T. & Kjeldsen, F. SuperQuant: A Data Processing Approach to Increase Quantitative Proteome Coverage. Anal Chem 87, 6319–6327, doi:10.1021/acs.analchem.5b01166 (2015).

23 Tyanova, S. et al. The Perseus computational platform for comprehensive analysis of (prote)omics data. Nat Methods 13, 731–740, doi:10.1038/nmeth.3901 (2016).

24 Ritchie, M. E. et al. limma powers differential expression analyses for RNA-sequencing and microarray studies. Nucleic Acids Res 43, e47, doi:10.1093/nar/gkv007 (2015).

25 Xia, J. G., Benner, M. J. & Hancock, R. E. W. NetworkAnalyst - integrative approaches for protein-protein interaction network analysis and visual exploration. Nucleic Acids Res 42, W167–W174, doi:10.1093/nar/gku443 (2014).

26 Shannon, P. et al. Cytoscape: A software environment for integrated models of biomolecular interaction networks. Genome Res 13, 2498–2504, doi:10.1101/gr.1239303 (2003).

